# Multiple domains of Matɑ1 transcription factor contribute to binding site specificity

**DOI:** 10.64898/2026.10.27.762204

**Authors:** Emily Knisely-Durham, Simone M. Giovanetti, Meru J. Sadhu

**Affiliations:** Center for Genomics and Data Science Research, National Human Genome Research Institute, National Institutes of Health, Bethesda, MD 20892, USA

## Abstract

Among homologous transcription factors with the same function, DNA-binding specificity can diverge. Matɑ1 is a high mobility group (HMG) box transcription factor in budding yeast whose *Saccharomyces cerevisiae* and *Candida albicans* homologs regulate ɑ-specific genes but bind distinct *cis*-regulatory sequences. To identify the protein features underlying this divergence, we generated structure-guided chimeras and measured their activity using species-specific reporters. Replacing any of the four Matɑ1 regions reduced activation of the *S. cerevisiae* reporter, while all chimeras retained activity on the *C. albicans* reporter. Thus, all four regions of Matɑ1 – not just the HMG box domain – are important for sequence specificity.

## Main text

High mobility group (HMG) box transcription factors regulate gene expression through interactions with DNA, but the features that determine their sequence specificity remain incompletely understood (Malarkey and Churchill 2012, Dragen et al., 2004). Sequences differences outside canonical DNA-binding domains can contribute to divergent transcription factor binding preferences (Gera et al., 2022). Understanding how different regions of a transcription factor contribute to sequence binding and gene regulation is important for understanding the evolution of these proteins and provide insights for computational predictions of genetic variants in disease models.

Matɑ1 is an HMG box transcription factor found in budding yeast that regulates mating type specific gene expression. In both *Saccharomyces cerevisiae* and *Candida albicans*, Matɑ1 activates ɑ-specific gene expression via interaction with Mcm1 (Baker et al., 2011). Although this regulatory function is conserved, the DNA sequences recognized by Matɑ1 are not; the Matɑ1 homolog from *S. cerevisiae* (*Sc*Matɑ1) prefers a CTATTG sequence and *Ca*Matɑ1 prefers a TCCTTT sequence. The adjacent Mcm1 recognition sequence at these genes, in contrast, is relatively conserved (Baker et al., 2011). The molecular basis for the Matɑ1 homologs’ difference in preferred binding sequence remains unclear.

To determine which regions of Matɑ1 contribute to species-specific regulatory activity, we divided *Sc*Matɑ1 and *Ca*Matɑ1 into four regions based on structural features as predicted by AlphaFold3 models of each Matɑ1 homolog complexed with its Mcm1 homodimer and bound to DNA, which defined, for both homologs, a putative Mcm1-interaction region with a long linker, an N-terminal flank, the HMG-box DNA-binding domain, and a C-terminal tail (Fig. 1A, B). We adapted the pair of reporter plasmids used by Baker et al. (2011), replacing the β-galactosidase with GFP while retaining either the *S. cerevisiae* or *C. albicans* Matɑ1-responsive promoter element upstream. As Baker et al. observed, *Sc*Matɑ1 drove expression from the promoter with the *S. cerevisiae* CTATTG sequence while *Ca*Matɑ1 did not. In contrast, we found that both Matɑ1 homologs could drive expression from the *C. albicans* TCCTTT sequence; while this differs from what Baker et al. observed, it allowed us to use the *C. albicans* promoter as a positive control for Matɑ1 functionality.

**Figure 1.**
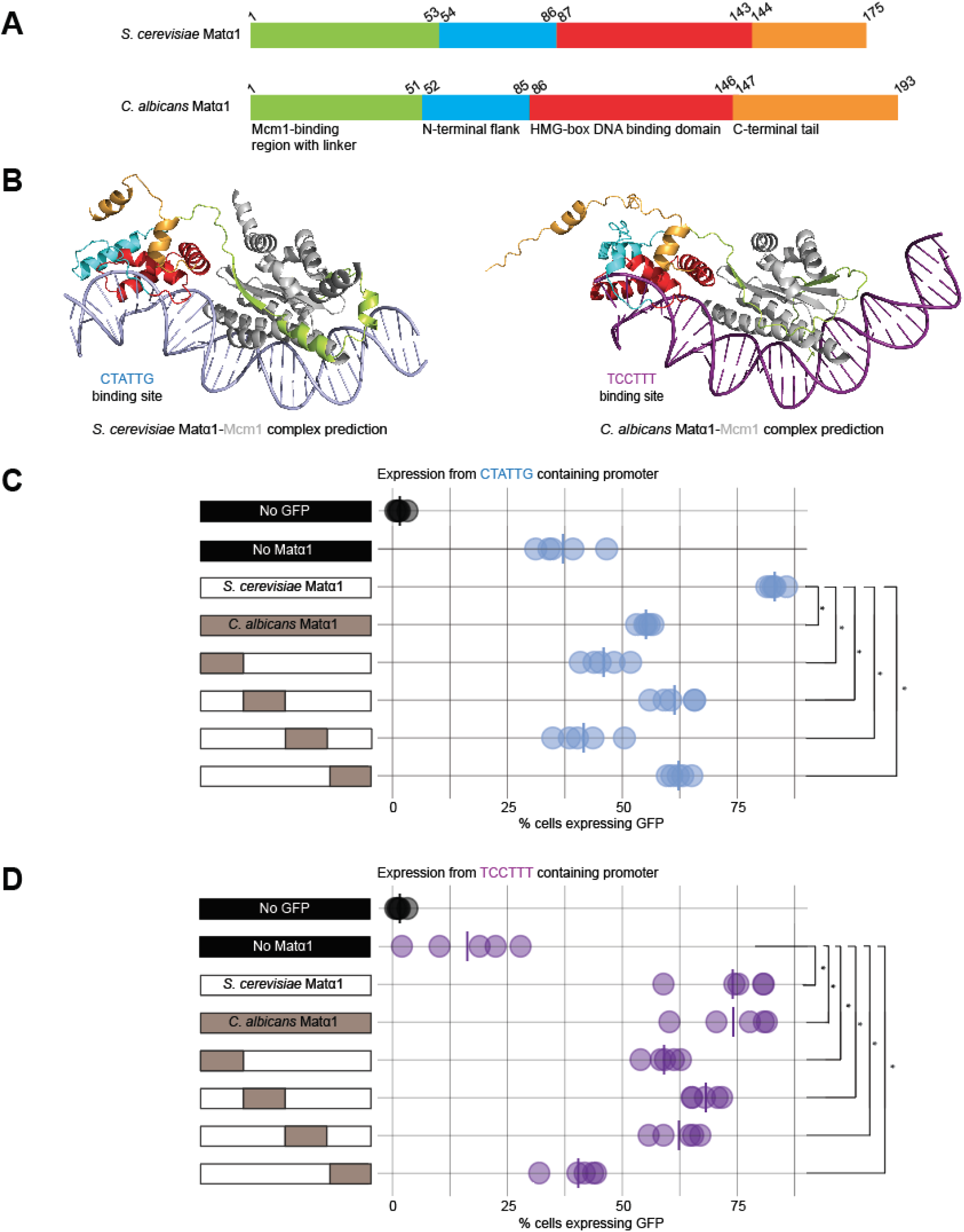
Matɑ1 regulates promoter sequence binding through all parts of the protein. A) Schematic of Matɑ1 domains important for regulating ɑ-cell type specific genes: predicted Mcm1-binding domain (green), N-terminal flank (cyan), HMG box domain (red), and C-terminal tail (orange). Each domain is colored according to the predicted structure in panel B. B) Predicted structure of Matɑ1-Mcm1 complex bound to DNA motifs for *S. cerevisiae* and *C. albicans* with AlphaFold3. For clarity, large unstructured N- and C-terminal tails of Mcm1 are not shown. C) Activation of the *S. cerevisiae* promoter sequence by full length and chimeric Matɑ1 proteins. D) Activation of the *C. albicans* promoter sequence by full length and chimeric Matɑ1 proteins. The same no-GFP negative control data are shown in panels C and D for reference.

To identify the regions of Matɑ1 responsible for these differences in regulatory activity, we generated chimeric proteins in which each of the four *S. cerevisiae* regions were individually replaced with the corresponding *C. albicans* sequence and measured activation of GFP reporters containing the *S. cerevisiae* CTATTG or *C. albicans* TCCTTT Matɑ1 recognition sequences.

Replacement of any of the four regions reduced activation from the *S. cerevisiae* CTATTG promoter relative to the full length *S. cerevisiae* Matɑ1, indicating that all four regions were required for activation. In contrast, just as both *Sc*Matɑ1 and *Ca*Matɑ1 activated the *C. albicans* TCCTTT promoter, we found that all four chimeras activated expression from that promoter above the level seen in the negative control fully lacking Matɑ1 (Fig. 1D), although the chimera sporting the *Ca*Matɑ1 C-terminal tail had diminished expression relative to the other chimeras. Thus, the chimeras’ inability to activate the *S. cerevisiae* CTATTG promoter was not due to lack of functional chimeric protein expression. Our results indicate that all regions contribute to the species-specific activity of Matɑ1.

Our results indicate that Matɑ1 sequence specificity is not determined solely by the canonical HMG-box domain. HMG proteins deform DNA upon binding, and sequence specificity can depend on the ability of a DNA sequence to accommodate the required conformation (King and Weiss, 1993, Dragan et al., 2004). Regions outside the HMG box may influence specificity by altering Matɑ1 conformation and DNA deformation, or they may contact DNA directly, as observed for regions flanking the HMG box in other HMG proteins (Love et al., 1995).

## Methods

### Plasmid construction

We obtained plasmids CB153 and CB154, containing the *Saccharomyces cerevisiae* and *Candida albicans* ɑ-specific regulatory sequences, respectively used by Baker et al. (2011). We replaced the β-galactosidase reporter with GFP. CB153 and CB154 were amplified with primers EKD5 and EKD44 to generate a linear vector fragment and GFP was amplified from MSp10 using primers EKD6 and EKD7. Fragments were assembled via Gibson assembly using a 5:1 molar ratio of insert to vector DNA, generating MSp758 (*S. cerevisiae* reporter) and MSp759 (*C. albicans* reporter).

Full length *S. cerevisiae* and *C. albicans* Matɑ1 expression plasmids and the four chimeric Matɑ1 plasmids were designed and synthesized by VectorBuilder. Chimera boundaries were guided by AlphaFold3 structural predictions. The four regions corresponded to residues 1-53, 54-86, 87-143, and 144-175 for *S. cerevisiae* and 1-51, 52-85, 86-146, and 147-193 of *C. albicans* Matɑ1. For each chimera, one *S. cerevisiae* region was replaced with the corresponding

*C. albicans* region in an otherwise *S. cerevisiae* Matɑ1 protein.

### Yeast strains and growth

The W303 Mata strain (MSy271) was transformed via a standard lithium acetate protocol (Gietz and Schiestl, 2007) with reporter plasmids alone or together with plasmids expressing full length or chimeric Matɑ1 proteins. Transformants were selected using the appropriate auxotrophic markers. Cultures were grown in CSM media lacking the appropriate amino acids at 30 C.

### Flow cytometry

Overnight cultures were diluted to OD600 = 0.1 and grown to mid-log phase (OD600 = 0.4). Cells were then diluted to OD600 = 0.1 in PBS in a 96-well plate and analyzed using a CytoFLEX S flow cytometer (Beckman Coulter). GFP fluorescence was measured for each sample. The percentage of GFP-positive cells was calculated for each biological replicate.

### Structural modeling

AlphaFold3 was used to generate structural predictions of Matɑ1-Mcm1-DNA complexes and guide the boundaries for chimera design. The *S. cerevisiae* model contained one *Sc*Matɑ1 protein, two *S. cerevisiae* Mcm1 proteins, and double-stranded DNA with the sequence 5′-TGTTCTATTGAAGGACTAATTAGGAAAGTTAAT-3′. The *C. albicans* model contained one *Ca*Matɑ1 protein, two *C. albicans* Mcm1 proteins, and double-stranded DNA with the sequence 5′-AAATTCCTTTCGCAGCCAAATTGGGTAACAAATAA-3′. Structures were visualized using PyMOL version 2.5.0.

### Data analysis and statistics

Flow cytometry data was analyzed in R using custom scripts. Statistical comparisons were performed using the Wilcoxon rank sum exact test. Code is available at https://github.com/SadhuLab/MATalpha1_Project.

### Plasmid table

|  |  |  |
|---|---|---|
| MSp756 | TEF1-*S. cerevisiae* Matα1-LEU2 | VectorBuilder |
| MSp757 | TEF1-*C. albicans* Matα1-LEU2 | VectorBuilder |
| MSp758 | *S. cerevisiae* α-specific promoter-GFP reporter, URA3 | Derived from CB153 |
| MSp759 | *C. albicans* α-specific promoter-GFP reporter, URA3 | Derived from CB154 |
| MSp760 | TEF1- *Sc*/*Ca* Matα1 chimera 1 (*Ca* 1-51 AA) in *Sc* Matα1 background, LEU2 | VectorBuilder |
| MSp761 | TEF1- *Sc*/*Ca* Matα1 chimera 2 (*Ca* 52-85 AA) in *Sc* Matα1 background, LEU2 | VectorBuilder |
| MSp762 | TEF1- *Sc*/*Ca* Matα1 chimera 3 (*Ca* 86-146 AA) in *Sc* Matα1 background, LEU2 | VectorBuilder |
| MSp763 | TEF1- *Sc*/*Ca* Matα1 chimera 4 (*Ca* 147-193 AA) in *Sc* Matα1 background, LEU2 | VectorBuilder |

### Strain table

| MSy271 | MATa ade2-1 ura3-1 his3-11,15 trp1-1 leu2-3,112 can1-100 |
|---|---|
| MSy336 | MSy271 with MSp758 |
| MSy337 | MSy271 with MSp758, MSp756 |
| MSy338 | MSy271 with MSp758, MSp757 |
| MSy339 | MSy271 with MSp758, MSp760 |
| MSy340 | MSy271 with MSp758, MSp761 |
| MSy341 | MSy271 with MSp758, MSp762 |
| MSy342 | MSy271 with MSp758, MSp763 |
| MSy343 | MSy271 with MSp759 |
| MSy344 | MSy271 with MSp759, MSp756 |
| MSy345 | MSy271 with MSp759, MSp757 |
| MSy346 | MSy271 with MSp759, MSp760 |
| MSy347 | MSy271 with MSp759, MSp761 |
| MSy348 | MSy271 with MSp759, MSp762 |
| MSy349 | MSy271 with MSp759, MSp763 |

### Oligonucleotide table

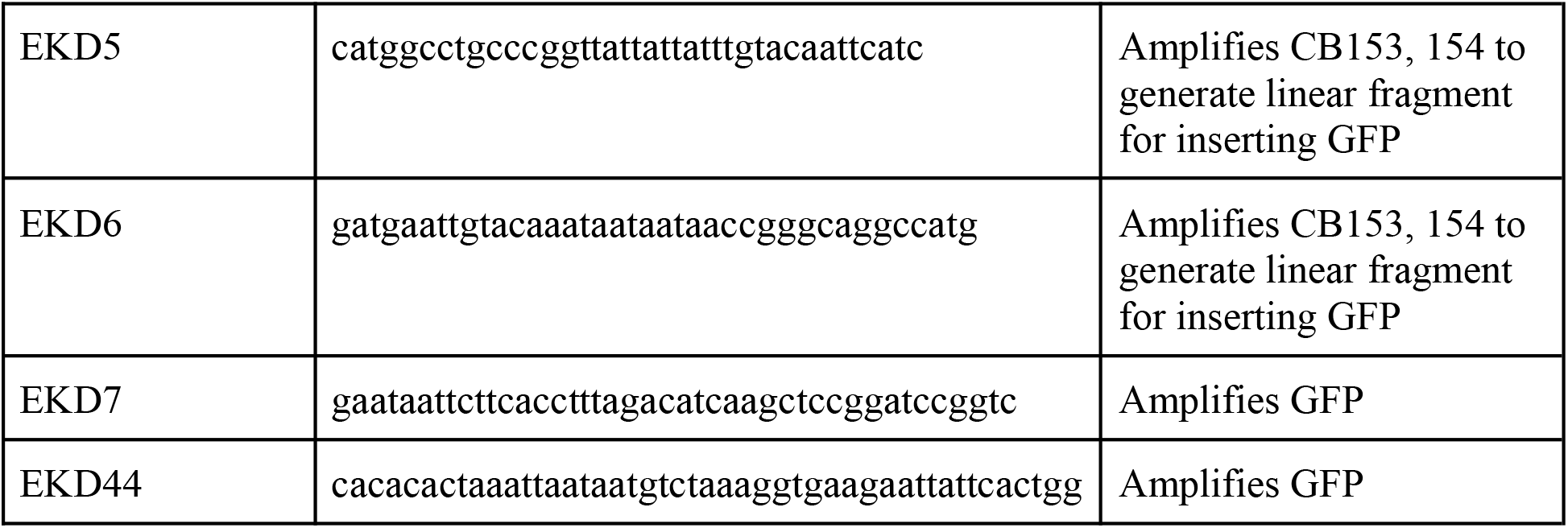

## Data availability

Code and data are available at https://github.com/SadhuLab/MATalpha1_Project.

## Acknowledgments

We thank Sandy Johnson for plasmids and Tom Dever for the parent yeast strain. We thank members of the Sadhu lab for helpful discussion. Flow cytometry was performed with the Flow Cytometry Core of the National Human Genome Research Institute. This work was supported by the Intramural Research Program of the National Human Genome Research Institute, NIH (1ZIAHG200401). The contributions of NIH authors are considered Works of the US Government. The findings and conclusions presented in this article are those of the authors and do not necessarily reflect the views of the NIH or the US Department of Health and Human Services.

## Author contributions

Conceptualization: M.J.S; Investigation and methodology: E.K-D., S.M.G., M.J.S.; .Writing: E.K-D., S.M.G., M.J.S

